# TITAN-BBB: Predicting BBB Permeability using Multi-Modal Deep-Learning Models

**DOI:** 10.64898/2026.02.15.706007

**Authors:** Gabriel Bianchin de Oliveira, Fahad Saeed

**Affiliations:** Knight Foundation School of Computing and Information Sciences, Florida International University, Miami, FL, USA

**Keywords:** Blood-Brain Barrier, Neural Networks, Transformers, CNN, Attention

## Abstract

Computational prediction of blood-brain barrier (BBB) permeability has emerged as a vital alternative to traditional experimental assays, which are often resource-intensive and low-throughput to meet the demands of early-stage drug discovery. While early machine learning approaches have shown promise, integration of traditional chemical descriptors with deep learning embeddings remains an underexplored frontier. In this paper, we introduce *TITAN-BBB*, a multi-modal deep-learning architecture that utilizes tabular, image, and text-based features and combines them using attention mechanisms. To evaluate, we aggregated multiple literature sources to create the largest BBB permeability dataset to date, enabling robust training for both classification and regression tasks. Our results demonstrate that TITAN-BBB achieves 86.5% of balanced accuracy on classification tasks and 0.436 of mean absolute error for regression, outperforming the state-of-the-art by 3.1 percentage points in balanced accuracy and reducing the regression error by 20%. Our approach also outperforms state-of-the-art models in both classification and regression performance, demonstrating the benefits of combining deep and domain-specific representations. The source code is publicly available at https://github.com/pcdslab/TITAN-BBB. The inference-ready model is hosted on Hugging Face at https://huggingface.co/SaeedLab/TITAN-BBB, and the aggregated BBB permeability datasets are available at https://huggingface.co/datasets/SaeedLab/BBBP.

## 1 Introduction

The Blood-Brain Barrier (BBB) is a layer of endothelial cells regulating substance exchange between the bloodstream and the Central Nervous System (CNS). Its function is to protect the CNS by ensuring essential nutrients to pass while blocking toxins and pathogens [1]. However, BBB presents a significant obstacle to drug development, effectively barring macromolecules and 98% of therapeutic small molecules from entering the brain [2].

Experimental evaluation of BBB permeability is challenging and costly. *In vitro* experiments often fail to replicate *in vivo* conditions, and in vivo studies are limited by invasiveness, high cost, and ethics [3]. These challenges make *in silico* models a valuable alternative, enabling scalable and efficient predictions, either qualitatively (i.e. if a compound is BBB permeable), or quantitatively (i.e. using logBB coefficient which is the ratio between the compound concentration in the brain and in the blood). Therefore, addressing this drug delivery challenge necessitates robust computational models capable of accurately predicting compound permeability across the BBB, and is an active area of research [4–7].

A variety of computational approaches have been proposed for BBB permeability prediction. Traditional machine learning models, such as LightGBM, Random Forest, and XGBoost [8, 9], rely on molecular descriptors from cheminformatics packages - limiting the learning to predefined features in a commercial database. On the other extreme, Graph Neural Networks (GNNs) [4], image-based, using convolutional neural networks, and text-based representations, such as mol2vec [10], learn embeddings directly from raw data represented as SMILES strings. Despite their individual strengths, the integration of domain knowledge with deep learning models has not yet been fully realized in the context of BBB permeability prediction, and remains underexplored [9].

In this paper, we introduce TITAN-BBB (**T**abular, **I**mage, and **T**ext combined via **A**ttention **N**etwork for **BBB** prediction), a multi-modal deep-learning architecture that integrates tabular, image, and text-based features derived from SMILES strings. Specifically, the model generates tabular descriptors using RDKit [11], image embeddings via ResNet50 [12], and text representations using ChemBERTa [13]. These multi-modal features are projected into a shared latent space and fused using a learnable attention mechanism. The resulting representation is processed by a prediction feedforward layer for classification or regression. To evaluate, we curated, and aggregated data from the literature [3,4,8,14–19]. To make the data ML-ready, we standardized the labels of the aggregated data resulting in the largest BBB permeability dataset to date - enabling robust training and evaluation on both classification and regression tasks. Our experimental results demonstrate that TITAN-BBB consistently surpasses state-of-the-art methods, achieving 86.5% of balanced accuracy on classification, and 0.436 of mean absolute error on regression tasks.

The rest of the paper is organized as follows. In Section 2, we describe the proposed method, the comparison approaches, the dataset, and the evaluation metrics. In Section 3, we evaluate and discuss the results for classification and regression tasks, the model interpretability and an ablation study. In Section 4, we present our conclusions and future work.

## 2 Materials and Methods

In this section, we present TITAN-BBB, the comparison models, the dataset, and the evaluation metrics.

### 2.1 Proposed Method

TITAN-BBB consists of three stages: multi-modal feature projection, attention-based fusion, and prediction. The pipeline of the model is presented in Figure 1.

**Figure 1:**
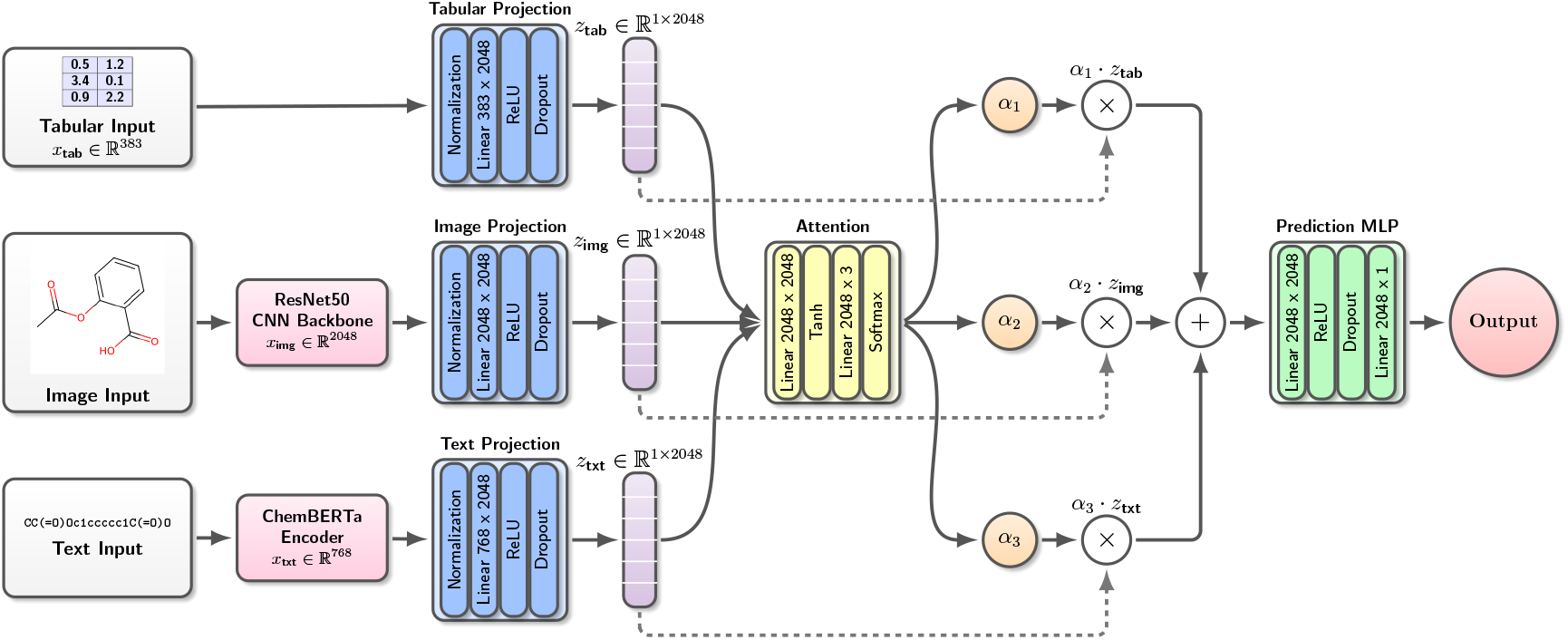
Overview of the TITAN-BBB architecture. The model takes SMILES strings as input to generate tabular descriptors, image embeddings via ResNet50, and text representations via ChemBERTa. Each modality is processed by a modality-specific projection block with identical architecture, composed of normalization, a linear layer, ReLU activation, and dropout, mapping heterogeneous input embeddings into a shared latent space of dimension 2048. Solid lines indicate the main data flow, while dashed lines indicate the routing of projected representations to the attention network for weight computation. A learnable attention network computes a scalar importance weight *α* for each modality, which are normalized via a softmax function. The projected modality vectors are scaled by their corresponding attention weights and combined through a weighted sum to form the final multimodal representation. This fused representation is passed to a feedforward head for classification or regression.

The model takes SMILES strings as raw input, from which tabular, image, and text representations are derived. For tabular features, we extract and concatenate a set of handcrafted descriptors composed of 217 RDKit 2D descriptors [11], which captures physicochemical properties such as molecular weight, polarity, and topological characteristics, and 166 MACCS keys [20], which are fixed-length binary fingerprints encoding the presence or absence of predefined chemical substructures. These descriptors were selected because they are well-established in the literature, have been utilized in BBB prediction methods [17, 21], and provide complementary information on global physicochemical properties and local structural motifs.

For image representation, SMILES strings are rendered as 2D molecular images using RDKit and processed by a frozen pretrained ResNet50 [12] network. To generate the embeddings, we employ the output from the final convolutional block of ResNet50. Regarding text embeddings, we utilize ChemBERTa-100M model [13] as a feature extractor. We employ the model with frozen weights, and obtain the final representation by computing the position-wise mean of embeddings for all valid tokens (excluding padding) to generate a single context vector per molecule.

To map the heterogeneous representations into a unified feature space, each modality is processed by a modality-specific projection block with identical architecture, as illustrated in Figure 1. Each projection block consists of a layer normalization followed by a linear layer that maps the input embedding (with modality-dependent dimensionality, i.e., 383 for tabular features, 768 for ChemBERTa embedding, and 2048 for ResNet50 embedding) into a common latent space of dimension 2048, followed by a ReLU activation and a dropout layer. The projection dimension was determined via grid search on the validation set, with 2048 yielding the best performance for both tasks.

Once the tabular, image, and text representations are projected into the shared feature space, we combine them using a learnable attention mechanism. This approach allows the model to dynamically weight the importance of each modality for every molecule. To determine the attention weights, we first compute an attention score *s*_*k*_ for each modality *k* ∈ {tab, img, txt} using a feed-forward network. Subsequently, these scores are normalized via a softmax function to obtain the final attention weights *α*_*k*_. Equation 1 presents the calculation of the attention weight *α* for each modality.

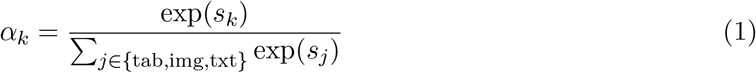

The final multi-modal representation *Z* is computed as the weighted sum of the input vectors, as shown in Equation 2.

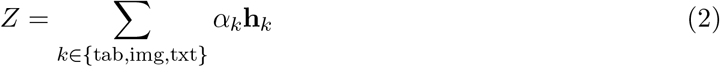

The fused vector *Z* is fed into a final prediction head. This stage is composed of a multi-layer feedforward network containing a linear layer with ReLU activation and dropout, followed by the final output layer.

We performed a grid search on the validation set to optimize hyperparameters (projection dimension and dropout). The optimal projection dimension was found to be 2048 for both tasks. Drop out rate of 0.1 for classification and 0.3 for regression tasks was used for experiments. For classification, the decision threshold was tuned to maximize performance considering the balanced accuracy score, with the optimal threshold determined at 0.7.

For training, we used a batch size of 32 over 1000 epochs with early stopping. The model was optimized using AdamW with a learning rate of 0.0001, employing weighted binary cross-entropy (for classification) or *L*_1_ loss (for regression).

### 2.2 Comparison with SOTA Methods

We compared our method against classical, image-based, fine-tuned transformer, and state-of-the-art approaches [4, 10, 17]. To ensure a fair and rigorous comparison, all models were retrained from scratch on our dataset (described in Section 2.3). Furthermore, we employed the validation set to optimize hyperparameters, ensuring they were evaluated at their optimal performance levels.

For classical models, we tested logistic regression (only for classification), random forest, Light-GBM, XGBoost, SVM (linear, polynomial, and RBF kernels), and k-NN, using RDKit [11] 2D descriptors, Morgan fingerprints, and MACCS keys as features. Hyperparameters and feature selection were optimized via grid search using the training set.

For image-based approaches, molecules were rendered as images using RDKit and processed with convolutional and attention-based networks, including ResNet18, ResNet50, ResNet101 [12], DenseNet121 [22], ViT [23], and Swin Transformers [24].

Considering transformer-based models, we fine-tuned ChemBERTa-100M, ChemBERTa-77M MLM, and ChemBERTa-77M MTR for SMILES sequence analyses.

Finally, for state-of-the-art methods, we used the available implementations of DeepBBBP [10]^1^, Deep-B3 [17]^2^, and MTGP [4]^3^. DeepBBBP combines convolutional and recurrent layers with mol2vec embeddings from SMILES. Deep-B3 aggregates features from a SMILES using LSTM, Morgan fingerprints, and image-based embeddings. Deep-B3 is only available for classification tasks. MTGP is a GNN-based method generating SMILES embeddings via masked pretraining, with the prediction performed using XGBoost.

### 2.3 Dataset

To construct our dataset, we aggregated and curated multiple sources from the literature [3, 4, 8, 14–19]. Because the sources varied in their annotation formats, we first standardized the labels. In some cases, compounds were annotated as BBB+ (crossing) or BBB- (non-crossing), while in others only a continuous logBB value was provided. To harmonize these representations for classification, we adopted the literature rule, i.e., compounds with logBB ≥ -1 were assigned to BBB+, and those with logBB *<* -1 to BBB-.

The initial aggregation of the sources yielded 16,603 unique compounds present in a single dataset and 8,112 duplicates found across at least two datasets. Among these duplicates, 7,703 carried consistent labels and were retained. To further refine the dataset, we transformed all SMILES into their canonical form and removed duplicates based on InChI keys, reducing the count from 24,199 to 9,299. Finally, compounds with ambiguous hydrogen configurations were filtered out, leaving a clean dataset of 9,262 compounds. The dataset is publicly available on our GitHub repository^4^ and on Hugging Face^5^.

For the classification task, we employed a scaffold split strategy to partition the dataset. By segregating compounds based on molecular structure, this approach ensures a more rigorous and realistic evaluation compared to a random split [21]. The data was divided into 80% for training, 10% for validation, and 10% for testing. Table 1 details the distribution of BBB+ and BBB-compounds across these subsets.

**Table 1:** Number of samples in each set for classification.

| Set Name | BBB+ | BBB- |
| --- | --- | --- |
| Training | 4,564 | 3,029 |
| Validation | 434 | 293 |
| Test | 638 | 304 |
| Total | 5,636 | 3,626 |

For the regression task, only compounds with experimentally available logBB values were considered. Since the regression subset was derived directly from the classification dataset, which was partitioned using a scaffold split, the train, validation, and test splits were preserved, ensuring that no structural leakage occurs between subsets. For compounds present in multiple sources, the logBB value was computed as the average across the sources, as the reported values exhibit low variance. This resulted in a reduced subset comprising 963 compounds for training, 84 for validation, and 100 for testing. The regression dataset is also publicly available on our repositories.

### 2.4 Evaluation Metrics

To evaluate classification performance, we used accuracy, balanced accuracy, specificity, sensitivity, precision, F1-score, and area under the receiver operating characteristic curve (AUC ROC). For the regression task, we evaluated model performance using mean absolute error (MAE), mean squared error (MSE), and root mean squared error (RMSE).

## 3 Results and Discussion

In this section, we compare TITAN-BBB against baselines and state-of-the-art models, analyze the model interpretability, and present an ablation study to assess the contribution of each modality.

### 3.1 Comparison with Baselines and SOTA

The outcomes of the proposed approach against baselines and comparison with the state-of-the-art are summarized in Table 2. In this table, for classification metrics, higher values are better, and for regression metrics, lower values are better.

**Table 2:** Performance of TITAN-BBB compared to baselines and state-of-the-art approaches on the test set. The top three results for each metric are highlighted: best in **blue**, second-best in **green**, and third-best in **orange**. Abbreviations: Acc. (Accuracy), Spe. (Specificity), Sen. (Sensitivity), Bal. Acc. (Balanced Accuracy), and Pre. (Precision).

| Model | Acc. | Spe. | Sen. | Bal. Acc. | Pre. | F1 | ROC AUC | MAE | MSE | RMSE |
| --- | --- | --- | --- | --- | --- | --- | --- | --- | --- | --- |
| TITAN-BBB | 0.857 | **0.888** | 0.842 | **0.865** | **0.940** | 0.888 | **0.935** | **0.436** | **0.399** | **0.632** |
| Logistic Regression | 0.838 | 0.789 | 0.861 | 0.825 | 0.896 | 0.878 | 0.825 | — | — | — |
| Random Forest | **0.880** | 0.760 | **0.937** | 0.849 | 0.891 | **0.914** | 0.849 | **0.454** | **0.403** | **0.635** |
| LightGBM | **0.878** | 0.806 | 0.912 | **0.859** | 0.908 | **0.910** | 0.859 | 0.485 | 0.436 | 0.660 |
| XGBoost | **0.879** | 0.799 | 0.917 | 0.858 | 0.906 | **0.911** | 0.858 | 0.507 | 0.470 | 0.685 |
| SVM Linear | 0.847 | 0.819 | 0.861 | 0.840 | 0.909 | 0.884 | 0.840 | 0.494 | **0.413** | **0.642** |
| SVM Polynomial | 0.872 | 0.750 | **0.929** | 0.840 | 0.886 | 0.907 | 0.840 | 0.462 | 0.419 | 0.647 |
| SVM RBF | 0.876 | **0.826** | 0.900 | **0.863** | **0.915** | 0.908 | 0.863 | **0.451** | 0.417 | 0.645 |
| k-NN | 0.864 | 0.766 | 0.911 | 0.839 | 0.891 | 0.901 | 0.839 | 0.514 | 0.444 | 0.666 |
| ResNet18 | 0.814 | 0.813 | 0.815 | 0.814 | 0.901 | 0.856 | 0.903 | 0.556 | 0.520 | 0.721 |
| ResNet50 | 0.839 | 0.806 | 0.854 | 0.830 | 0.902 | 0.878 | **0.915** | 0.603 | 0.596 | 0.772 |
| ResNet101 | 0.773 | **0.875** | 0.724 | 0.800 | **0.924** | 0.812 | 0.901 | 0.547 | 0.540 | 0.735 |
| DenseNet121 | 0.839 | 0.786 | 0.864 | 0.825 | 0.894 | 0.879 | 0.911 | 0.578 | 0.566 | 0.752 |
| ViT | 0.832 | 0.720 | 0.886 | 0.803 | 0.869 | 0.877 | 0.888 | 0.567 | 0.565 | 0.751 |
| Swin Transformers | 0.798 | 0.822 | 0.787 | 0.805 | 0.903 | 0.841 | 0.897 | 0.581 | 0.569 | 0.754 |
| ChemBERTa-100M | 0.848 | 0.753 | 0.893 | 0.823 | 0.884 | 0.889 | 0.909 | 0.588 | 0.549 | 0.741 |
| ChemBERTa-77M MLM | 0.817 | 0.661 | 0.892 | 0.777 | 0.847 | 0.869 | 0.842 | 0.608 | 0.598 | 0.773 |
| ChemBERTa-77M MTR | 0.850 | 0.717 | 0.914 | 0.815 | 0.871 | 0.892 | 0.895 | 0.623 | 0.615 | 0.784 |
| DeepBBBP | 0.677 | 0.000 | **1.000** | 0.500 | 0.677 | 0.808 | 0.409 | 0.602 | 0.599 | 0.774 |
| Deep-B3 | 0.837 | 0.747 | 0.879 | 0.813 | 0.879 | 0.879 | 0.907 | — | — | — |
| MTGP | 0.861 | 0.757 | 0.911 | 0.834 | 0.887 | 0.899 | **0.922** | 0.548 | 0.472 | 0.687 |

For the classification task, the proposed method achieved superior performance, obtaining the best results in four of seven metrics (88.8% of specificity, 86.5% of balanced accuracy, 94.0% of precision, and 92.2% of ROC AUC) as compared to the other methods. Balanced accuracy is the most significant metric for this task, as it provides a balanced calibration between specificity and sensitivity, with TITAN-BBB achieving the highest score on this metric. These results consistently indicate that the approach is competitive and surpasses most baselines across different evaluation criteria.

Furthermore, for the regression task, TITAN-BBB obtained the best results across all metrics, including 0.436 of MAE, 0.399 of MSE, and 0.632 of RMSE. Among these metrics, MAE is the most relevant for this task, as it directly reflects the average absolute prediction error. The lower MAE indicates that TITAN-BBB provides more accurate estimates of BBB permeability compared to all other comparison methods.

Finally, when compared against state-of-the-art methods, the proposed approach consistently achieved superior results across all scenarios. Specifically, TITAN-BBB outperformed the previous state of the art by 3.1 percentage points in balanced accuracy for classification and reduced the mean absolute error by approximately 20% for regression. These results highlight the effectiveness of the proposed model for BBB permeability prediction across both tasks.

To evaluate the robustness of the proposed approach, we assessed selected models under two additional scaffold split ratios: 70/15/15 and 60/20/20 for training, validation, and test sets, respectively. The selected models correspond to the top-performing methods on the original 80/10/10 split. Table 3 presents the results. TITAN-BBB consistently obtained the lowest MAE across all splits, confirming its superiority in regression tasks. For classification, a performance decrease is observed with reduced training data, which is consistent across all evaluated methods and reflects the increased difficulty imposed by the scaffold split strategy. Models with a higher number of parameters, such as TITAN-BBB, are more sensitive to reductions in training set size.

**Table 3:**
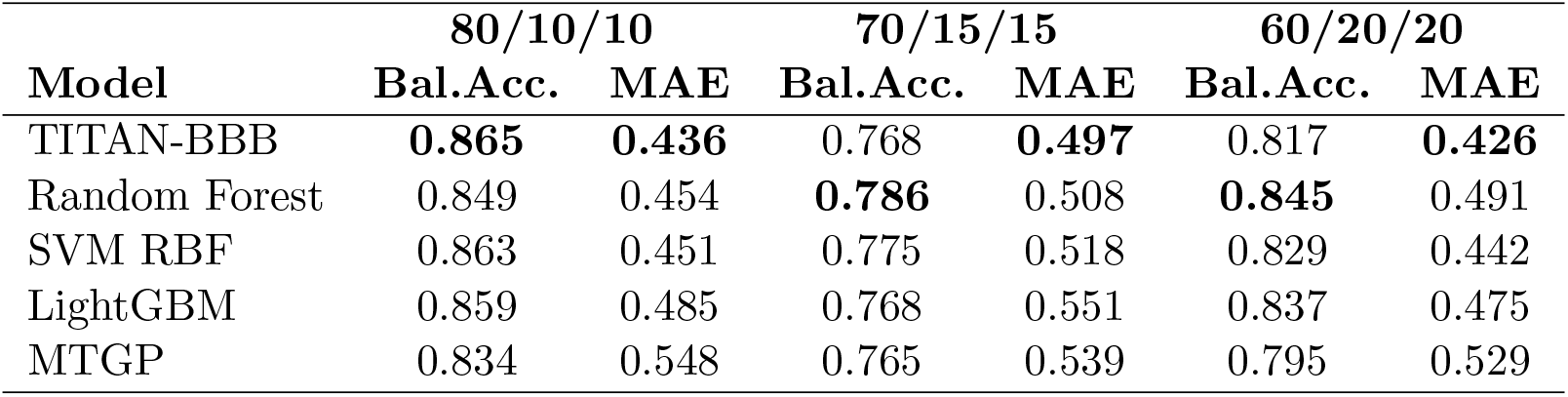
Robustness evaluation of TITAN-BBB and selected baselines across scaffold split ratios. Results report Balanced Accuracy (Bal. Acc.) for classification and MAE for regression. The best results are in **bold**.

### 3.2 Interpretability

To investigate the contribution of each component, we performed an interpretability analysis based on the attention value of each modality. Figure 2 presents the average importance value of each modality across the test set. For both tasks, the tabular features demonstrated the highest importance, consistently exhibiting an average attention value above 60%. Following this, the textual features were the second most important, contributing approximately 20% in both tasks. Finally, the image modality showed the lowest importance.

**Figure 2:**
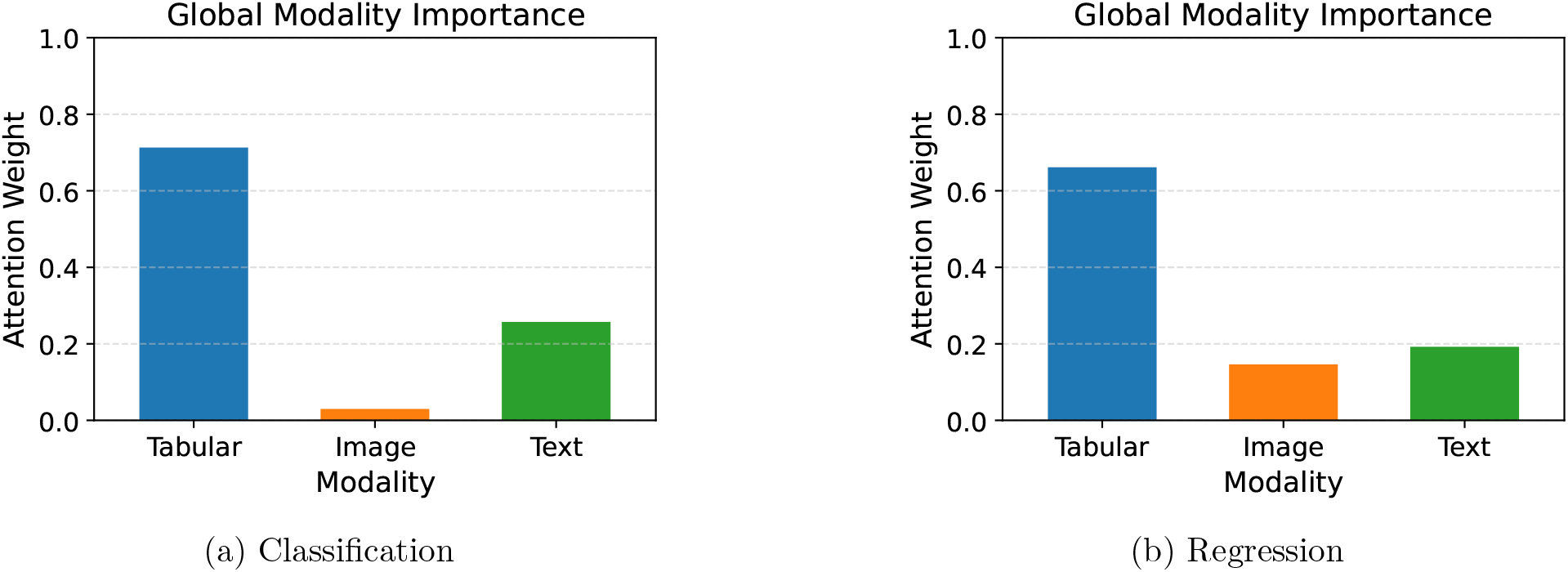
Attention weights for each modality considering the classification (a) and regression (b)tasks. For both tasks, the tabular modality obtained the highest attention values, followed sequentially by the textual and image modalities.

Since the tabular data are composed of 2D descriptors and MACCS keys, we calculated the importance of each feature set using Integrated Gradients [25]. The graphical results for both classification and regression tasks are illustrated in Figure 3. The analysis shows that both categories of tabular features exhibited nearly similar mean importance for the test set, with the MACCS keys demonstrating a slight predominance.

**Figure 3:**
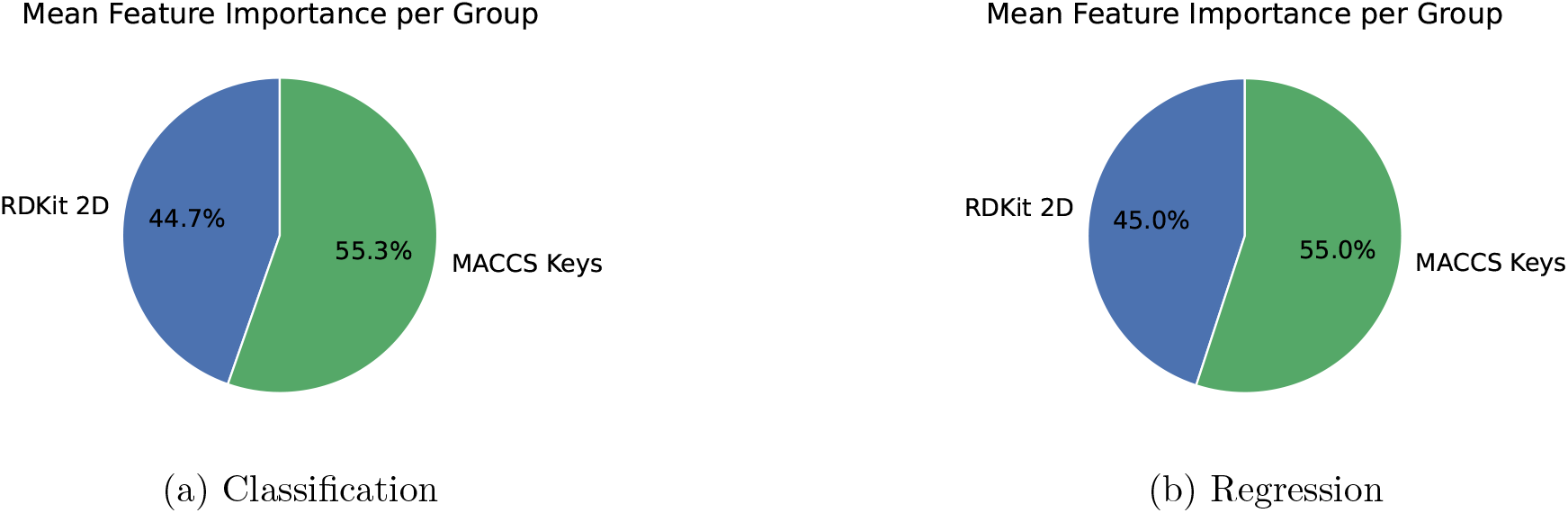
Integrated Gradients of tabular features considering the classification (a) and regression (b)tasks. For both tasks, 2D descriptors and MACCS keys showed similar importance, with MACCS keys demonstrating a slightly higher importance.

To validate the quality of the multi-modal representations, we analyzed the features generated for the nicotine molecule, which is included in the test set. This analysis is presented in Figure 4. For the image modality, we applied EigenCAM [26] to produce a heatmap, which indicates that the embeddings are primarily driven by the aromatic regions and the central linkage between the rings. This suggests that the image embeddings are able to capture global molecular topology, despite having been pretrained on natural images.

**Figure 4:**
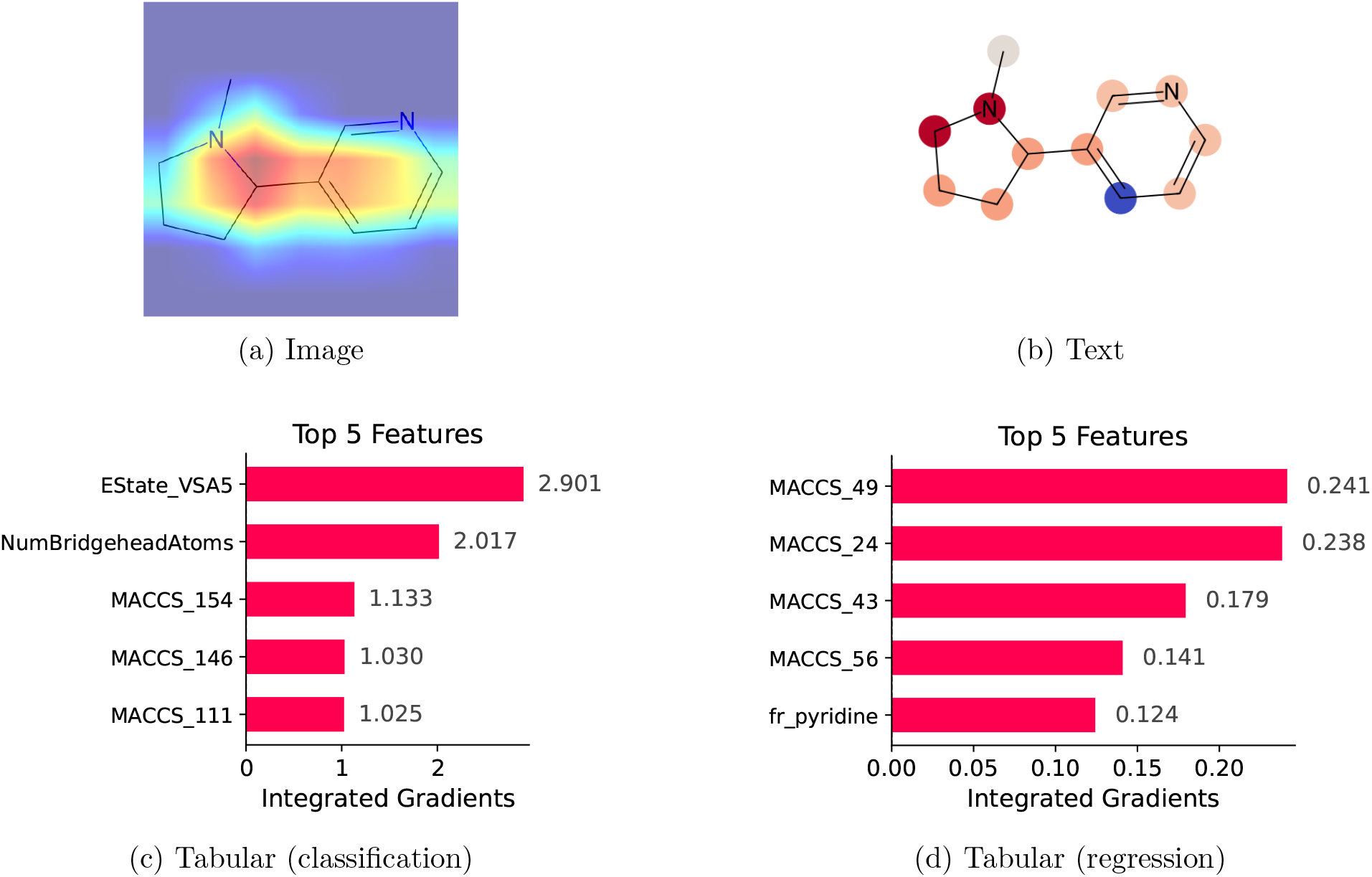
Interpretability of nicotine molecule representation considering image (a), text (b), tabular for classification (c), and tabular for regression (d). Image and text embeddings are the same for classification and regression tasks.

For the text modality, where the final representation is the mean of token embeddings, we used SHAP [27] to assess the contribution of each token to the final textual features. Token importance was subsequently mapped to atom importance. In the figure, red indicates higher relevance of a given atom, whereas blue denotes lower importance. The SHAP values show that the aggregated embedding is strongly influenced by the nitrogen atom. Based on this result, we conclude that the frozen ChemBERTa prioritizes functional groups over generic structural tokens for nicotine molecules.

Finally, for the tabular modality, the Integrated Gradients analysis confirms that explicit descriptors, such as MACCS keys topological patterns, regional electronic properties (EState VSA5), structural rigidity (NumBridgeheadAtoms), and functional motifs (fr pyridine) were the decisive factors for the final prediction.

These analyses indicate that each modality provides a distinct and complementary view of the molecule. While the image representation captures the global topology, the text embedding focuses on local semantic features and specific reactive atoms, and the tabular features pay attention to explicit physicochemical properties. The resulting model TITAN-BBB, therefore, learns from multimodal data and performs predictions better than single modality models.

### 3.3 Ablation Study

To further assess the contribution of each component, we conducted an ablation study. During this evaluation, we retrained each TITAN-BBB variant from scratch to evaluate the predictive capability of each modality. The results are presented in Table 4. We observed that while the tabular modality is a strong individual predictor (83.4% of balanced accuracy and 0.453 of MAE), the deep learning modalities also demonstrate independent predictive capabilities when trained in isolation, achieving 80.0% and 77.4% of balanced accuracy and 0.581 and 0.580 of MAE, respectively. Regarding single-modality exclusion, the exclusion of tabular representations causes the most significant performance drop. However, excluding image or text embeddings also degrades performance, confirming that TITAN-BBB relies on the complementarity between tabular, image, and textual representations to achieve state-of-the-art performance.

**Table 4:** Ablation study of TITAN-BBB. Each configuration was trained from scratch to isolate the contribution of specific modalities. The results demonstrate the performance degradation when specific modalities are excluded compared to the original TITAN-BBB, highlighting the dependency on multi-modal integration.

| Configuration | Bal. Acc. | MAE |
| --- | --- | --- |
| TITAN-BBB | 0.865 | 0.436 |
| tabular | 0.834 | 0.453 |
| image | 0.800 | 0.581 |
| text | 0.774 | 0.580 |
| image and text | 0.816 | 0.599 |
| tabular and text | 0.854 | 0.465 |
| tabular and image | 0.838 | 0.454 |

## 4 Conclusion

In this work, we introduce TITAN-BBB, a multi-modal deep-learning model for BBB prediction using tabular, image, and text representations. Our approach outperforms state-of-the-art models, achieving 86.5% of balanced accuracy and 0.436 of MAE for classification and regression tasks, respectively. As part of this work, we have downloaded, collected, labelled, and integrated a curated BBB dataset that can be used as training, validation, and testing data and may serve as a benchmark for both regression and classification tasks. To the best of our knowledge, this is the largest dataset available for BBB permeability prediction in both settings, with 9,262 compounds for classification and 1,147 compounds for regression. The data set is being made available at https://huggingface.co/datasets/SaeedLab/BBBP.

In addition to the overall performance metrics, our interpretability and ablation studies also demonstrate that each modality of the data is learning something useful for BBB prediction, resulting in a model that is better than a single modality model.

## Acknowledgment

Research reported in this publication was supported by NIGMS of the National Institutes of Health under award number R35GM153434. The content is solely the responsibility of the authors and does not necessarily represent the official views of the National Institutes of Health.

## Footnotes

1 https://github.com/dasdibye/bbbpDL

2 https://github.com/GreatChenLab/Deep-B3

3 https://github.com/mldlproject/2025-MGTP

4 https://github.com/pcdslab/TITAN-BBB

5 https://huggingface.co/datasets/SaeedLab/BBBP

